# First report of geophagy by chimpanzees in Comoé National Park, Ivory Coast

**DOI:** 10.1101/2023.07.25.550531

**Authors:** Juan Lapuente, K. Eduard Linsenmair

## Abstract

West African chimpanzees (*Pan troglodytes verus*) are critically endangered, therefore knowing their ecological needs is necessary to implement proper conservation measures, especially in the face of climatic change. We report here the consumption of soil by wild chimpanzees living in Comoé National Park, Ivory Coast. We confirmed through camera-trap recording that chimpanzees of all ages and both sexes consumed it for several years at least in one community and more occasionally in a second one, aided by leaf-sponge tools. Our results suggest that these chimpanzees living in a savanna-forest mosaic may need minerals supplement, including sodium or/and clay to aid digestion, mainly during the dry season.

## 1 INTRODUCTION

Wild chimpanzees are known to eat soil (geophagy) across Africa. This behavior was already described by Goodall in Gombe andNishida and Uehara in Mahale in the decade of 1960 (Pebsworth et al., 2019) and it has also been found in West African chimpanzee in Fongoli, Senegal (Gašperšič and Pruetz, 2011). In many cases, including the previous, the apes have been described to also consume the soil from termite mounds, mainly of genera Cubitermes and Macrotermes. It has been suggested that this ingestion of soil helps the chimpanzees and other primates to obtain minerals or to calm gut ailments (Krishnamany and Mahaney, 2000; Ketch et al. 2001) or to increase medical properties of plants they consume (Klein et al. 2007). In other cases, chimpanzees have consumed directly the soil from the ground or cliffs, frequently exposed by the erosion or the activity of other animals in what are called salt-licks. Popularly, the salt-licks are believed to attract the animals in quest of sodium, but analysis of some geophagy sites found that animals may also consume sodium poor soils, for instance, Reynolds et al. (2015, 2019) and Pebsworth et al. (2018,2019) consider that the iron and aaluminium rich soil consumed by chimpanzees in Budongo Forest (Uganda) is mainly useful to neutralize the tannins, alcaloids and phenolics present in some vegetable foods, while Mahaney et al. (2005) suggested that chimpanzees in Kibale Forest, Uganda, eat the soil to supplement their Iron poor diet.

Even though chimpanzees are thought to obtain most of the minerals they need from their diet, they may not obtain enough sodium, needing an extra supply from soil, insects or some specific plants (O’Malley et al. 2014, Pebsworth et al. 2018, Reynolds et al .2019), the ones living in savanna habitats might need an extra supply of this last mineral, since they may lose more due to increased transpiration, especially during the dry season, which could lead them to search for additional sources of sodium during the dry season, such as sodium rich soil. We wanted to know if Comoé chimpanzees (*Pan troglodytes verus*) consumed the soil from the so called salt-licks, how often they did it, if all individuals in a community did it and if they did it seasonally or all the year long.

## 2 METHODS

Comoé National Park (CNP) is the biggest protected area in Ivory Coast, with 11500 km2. Chimpanzees live in a savanna-forest mosaic with around 13 % of dry and semideciduous forest cover. Annual precipitation is around 1090 mm (925 mm during the study period) and mean annual temperature of 27° C, with a dry season from mid October to mid April and a wet season the rest of the year. Precipitation is mostly concentrated in the wet season, with peaks in May and September, while the months of December and January virtually lack precipitation. The hottest months are February and March, with mean maximum temperatures above 35º C, while, during rainy season, mean maximum temperature is below 26º C. December and January are also the coldest months because of the harmattan cool winds. (Lapuente, 2021).

To answer our questions, we searched for geophagy sites (salt-licks) within the home-ranges of four of our chimpanzee study communities in CNP, two in the savanna, less than 100 m away from the forest edge, close enough to the forest to be visited by chimpanzees, and four within the forest itself. We placed camera traps at these six sites, a technique that had previously been used to monitor soil consumption by other authors (Pebsworth et al. 2018, 2019). Comoé chimpanzees are non-habituated, therefore, we study them mainly through indirect signs and camera trapping. One of the geophagy sites in the savanna and three in the forest were in cliffs composed of marlstone (a multilayered, soft, clay-rich stone), which had been dug by a variety of animals when eating the soil. The other two sites were on flat ground, in red, clay-rich (> 80 % clay) lateritic soil, also dug by animals. To determine the soil texture, we powdered and sieved four samples of 50 g of soil from every site and decanted them in a jar with water for 48 h, measuring the percentages of the clay, silt and sand fractions with a milimeter to determine proportion of clay, sand and silt in the sample. We used the soil classification used by US agriculture department. We tested the presence of carbonate in the samples by dropping one drop of diluted (10 %) hydrochloric acid and waiting for bubbles of CO2 to emerge.

We continuously monitored with camera traps the four geophagy sites where we recorded no visits of chimpanzees for six months and the two where we did record visits of chimpanzees for 24 months, including therefore, for these, two full dry seasons and two full wet seasons. The camera traps were Bushnell HD and Browning SPEC OPS, programmed to record 60 second videos and working permanently (24 h/day) for the study period. We used the age classes defined by Sugiyama (1999) adult: > 11 years, adolescent: 8–11 years, juvenile: 4–7 years, infant: < 4 years

## 3 RESULTS

### 3.1 Soil analysis

The analysis of the soil showed that the marl extracted from the cliffs contained much less clay than the red latheritic soil in flat ground (Table 1) On the other hand, we confirmed the presence of carbonate in the marl samples, which was probably calcium carbonate (calcite) from the vigorous reaction. The fraction identified as silt in the marl samples was therefore formed mainly by calcite or limestone.

**Table 1.** Clay, sand and silt fractions percentages obtained with the jar test (see methods) for the samples collected in four geophagy sites (salt-licks) and two control regular soil samples collected close to the two sites where chimpanzees consumed soil and soil filled water with leaf-sponges. (Means ± SD of four samples per site)

| SAMPLE | Clay % | Sand % | Silt % | Classification of soil |
| --- | --- | --- | --- | --- |
| 1 Marl cliff in forest where Odyssey chimpanzees visited the most | 40.75 ± 1.26 | 16 ± 0.81 | 43.25 ± 0.96 | Silty clay |
| 2 Marl cliff in savanna, no chimpanzee visit. Many visits of baboons and other animals | 39.5 ± 17.14 | 22.5 ± 9.76 | 38 ± 16.63 | Silty clay loam |
| 3 Clay-rich, flat ground, where Achean chimpanzees drank with leaf-sponges | 81.75 ± 28.9 | 3.25 ± 3.03 | 15 ± 1.01 | Clay |
| 4 Clay-rich, flat ground, no chimpanzee visit. Many visits of other animals | 84.75 ± 25 | 2.5 ± 0.55 | 12.75 ± 5.31 | Clay |
| 5 Control regular soil close to (1) marl cliff in forest where Odyssey chimpanzees visited the most | 43.5 ± 8.29 | 12.5 ± 5.46 | 44 ± 17.36 | Silty clay |
| 6 Control regular soil close to (3) where Achean chimpanzees drank with leaf-sponges | 81.25 ± 32.63 | 6.75 ± 1.01 | 12 ± 2.5 | Clay |

### 3.2 Camera-trapping

We recorded the chimpanzees of three different communities visiting three of the geophagy sites (salt-licks), two on cliffs in the forest and one on flat-ground in the forest. We never recorded the chimpanzees visiting the salt-licks in the savanna. In the salt-lick on flat-ground, we did not record the chimpanzees eating the soil, but we recorded two adolescent individuals (one male and one female) using leaf-sponges to drink clay rich water accumulated in the holes of the salt-lick after a rain-storm. We previously had found four more leaf-sponges left on the ground in this salt-lick. In one of the forest cliffs, two chimpanzees just passed by, without consuming the soil. In the other forest cliff, we recorded 21 of the chimpanzees of Odyssey community, of all sex and age classes, coming repeatedly to consume the soil (clay-rich marlstone) that they scraped with fingernails or with teeth, remaining for an average of 1:55 minutes (Table 2) and occasionally napping at the site.

**Table 2.** Number of visits to the salt lick by the different individuals of the Odyssey community recorded, distinguishing between wet and dry season and visit in which they consumed soil.

| CHIMPANZEE | SEX/AGE | NUMBER OF VISITS RECORDED DRY SEASON | NUMBER OF VISITS RECORDED WET SEASON | NUMBER OF VISITS WHEN EATING SOIL | MEAN TIME EATING SOIL RECORDED PER VISIT (MINUTES) |
| --- | --- | --- | --- | --- | --- |
| ITHACA | ADOLESCENT F | 3 | 1 | 3 | 01:43 |
| LAERTES | ADOLESCENT M | 4 | 2 | 6 | 02:04 |
| ORESTES | ADOLESCENT M | 1 | 1 | 2 | 01:50 |
| ANTICLEA | ADULT F | 4 | 1 | 2 | 00:40 |
| CIRCE | ADULT F | 5 | 3 | 5 | 01:01 |
| EURICLEA | ADULT F | 3 |  | 2 | 06:35 |
| KTIMENE | ADULT F | 4 | 2 | 5 | 02:20 |
| MELANTEA | ADULT F | 4 |  | 3 | 01:20 |
| PENELOPE | ADULT F | 1 |  | 0 | 00:00 |
| POLIFEMO | ADULT M | 2 |  | 2 | 01:45 |
| CIRCE'S INFANT | INFANT | 6 | 3 | 9 | 01:37 |
| EURICLEA'S INFANT | INFANT | 2 |  | 2 | 04:20 |
| KTIMENE'S INFANT | INFANT | 3 |  | 2 | 01:10 |
| MELANTEA'S INFANT | INFANT | 2 |  | 0 | 00:00 |
| UNID. INFANT 1 | INFANT | 2 |  | 2 | 02:10 |
| UNID. INFANT 2 | INFANT | 2 |  | 2 | 02:00 |
| UNID. JUVENILE | JUVENILE | 3 |  | 2 | 00:35 |
| CALYPSO | JUVENILE F | 2 |  | 2 | 04:05 |
| INO | JUVENILE F | 5 | 3 | 8 | 02:19 |
| NAUSICAA | JUVENILE F | 4 | 1 | 4 | 00:54 |
| EURILOCO | JUVENILE M | 7 |  | 5 | 01:48 |
| **Total Result** |  | **69** | **17** | **68** | **01:55** |

Pooling together all the 86 visits to the salt lick of chimpanzees of all sex/ages of Odyssey community, we found that 69 were during the dry season and 17 during the wet season, which gives an 80 % of visits during the dry season. The chimpanzees, however, did not consume soil in every visit, only in 68 of the occasions, 54 in the dry season and 14 in the wet season, this is 79.4 % of the soil consumption episodes in the dry season.

Apart from the chimpanzees, we recorded three other primate species eating the soil in the marl cliff in the forest, Lowe’s monkey (*Cercopithecus lowei*) lesser puty nosed monkey (*Cercopithecus petaurista*) and olive baboon (*Papio anubis*), which was the primate species that visited the cliffs (both savanna and forest) most frequently. Apart from the primates, only herbivores visited the salt-licks and ate the soil (both in savanna and forest), including forest elephant (*Loxodonta africana*), buffalo (*Syncerus caffer*), giant hog (*Hylochoerus meinertzhageni*) and many others (table I in supplementary materials), being the most frequent consumers the kob (*Kobus kob*) and hartebeest (*Alcelaphus buselaphus*) in the savanna and the bushbuck (*Tragelaphus scriptus*) and Maxwell’s duiker (*Philantomba maxwelli*) in the forest.

## DISCUSSION

We confirmed that Comoé chimpanzees eat the soil of a marl cliff exposed by the erosion and the soil consumption by other animals, while we could not confirm if they consume the redclay in other geophagy sites of their home range, except for two adolescent individuals drinking the clay-enriched water with leaf-sponges in one of the sites. We confirmed that individuals of all sex/age classes in the Odyssey community visited the marl cliff and 19 of them where recorded to consume the soil. The fact that they visited this geophagy site mainly in the dry season (80 % of the visits) suggests that they need the soil more frequently in this period of the year. The fact that the chimpanzees consumed often the marl and we never recorded them consuming the red clay suggests that they obtain something else than simply clay from this soil, possibly minerals, but we need further analysis to confirm this. Pebsworth at al. (2018) cites Goodall already speculating back in 1968 at chimpanzee’s eating the soil to obtain extra salt, but she did not test this hypothesis. Other authors that more recently analyzed the soils consumed by chimpanzees in Budongo, Uganda, found that they were richer in aaluminium (Reynolds et al. 2015, 2019) or Iron (Mahaney et al. 2005, Pebsworth et al. 2019) than in sodium, suggesting that the chimpanzees supplemented their diets for these minerals or consumed the clay in the soil mainly to neutralize the toxicity of tannins, alcaloids and phenolics rich foods. We have not analyzed the soil content for minerals in this study, however, we did find that the marlstone consumed by the chimpanzees contained calcium carbonate and less clay than the samples obtained from flat ground geophagy sites not visited by chimpanzees, suggesting that they did not consume the soil just to obtain clay, but to obtain probably minerals. Only the salt-licks in cliffs of marlstone were especially rich in limestone, while other so called salt-licks by the locals (“salines” in French) were just rich in clay. The two adolescents of the Achean community that we recorded drinking clay-enriched water may have been just drinking from an accesible pool or trying to obtain other minerals from the clay.

Nevertheless, in the case of chimpanzees of Odyssey community, they visited the marlstone cliff in 86 occasions consuming the soil mainly in the dry season, which suggests that they did look for minerals and they looked for them more frequently when they had higher need, in the dry season. If they just looked for clay to neutralize toxics in food consumed during the dry season, they could have consumed the much more clay-rich soil from the flat ground geophagy sites, but they did not. However, the fact that the chimpanzees consumed less soil during the wet season could be related to the better sources of minerals that they consume during this period, mainly the social insects, ants and termites, which have been pointed as an important source for these and other nutrients (O’Malley et al. 2014). We recorded Odyssey’s individuals of all sex/age classes visiting repeatedly the salt-lick during two years, which suggests that this behavior may be customary in this chimpanzee community of Comoé. The fact that we could not record chimpanzees in the other five study communities that we follow in Comoé consuming the soil in the same manner yet, could be because we have not found their main geophagy sites yet or because they have other extra sources of minerals that we could not detect so far, although the habitat conditions are very similar for all the study communities, except Theogony, which has all its home-range covered by forest.

Comoé chimpanzees also consume sizable ammounts of Ceiba pentandra inner bark during the peak of the rainy season (Lapuente et al. 2020) when other food sources are scarce. This has been interpreted as a possible fallback food, but could also be a supplementary source of minerals during this period of food scarcity, when Comoé chimpanzees also consume less soil.

The fact that other animals consumed the soil both of the sodium rich marl cliffs and the sodium poor clay salt-licks could be because they look for sodium in the marl and other minerals in the clay. Interestingly, the animals that visited most frequently both types of salt-licks feed on leaves (bushbucks) and unripe fruits dropped by monkeys (duikers) which suggests that they could be trying to fight the toxicity of their foods. The primates that visited the salt-licks most frequently, however, the baboons, have a varied, fruit rich diet, similar to that of chimpanzees, although with a higher proportion of grass seeds and bark. We may speculate that baboons need more clay for their tannin richer diet and more sodium for the higher transpiration that their longer time in savanna could produce, but more research is needed to validate these speculations. Similarly, more research is needed to confirm if all the chimpanzee communities in Comoé display this behavior and to ascertain if they could have other motivations to consume soil. We also would need to wait for the habituation of some of these communities to find out if they also consume soil from other sources, such as termite mounds, which has never been recorded with our camera traps to the date.

**Figure 1.**
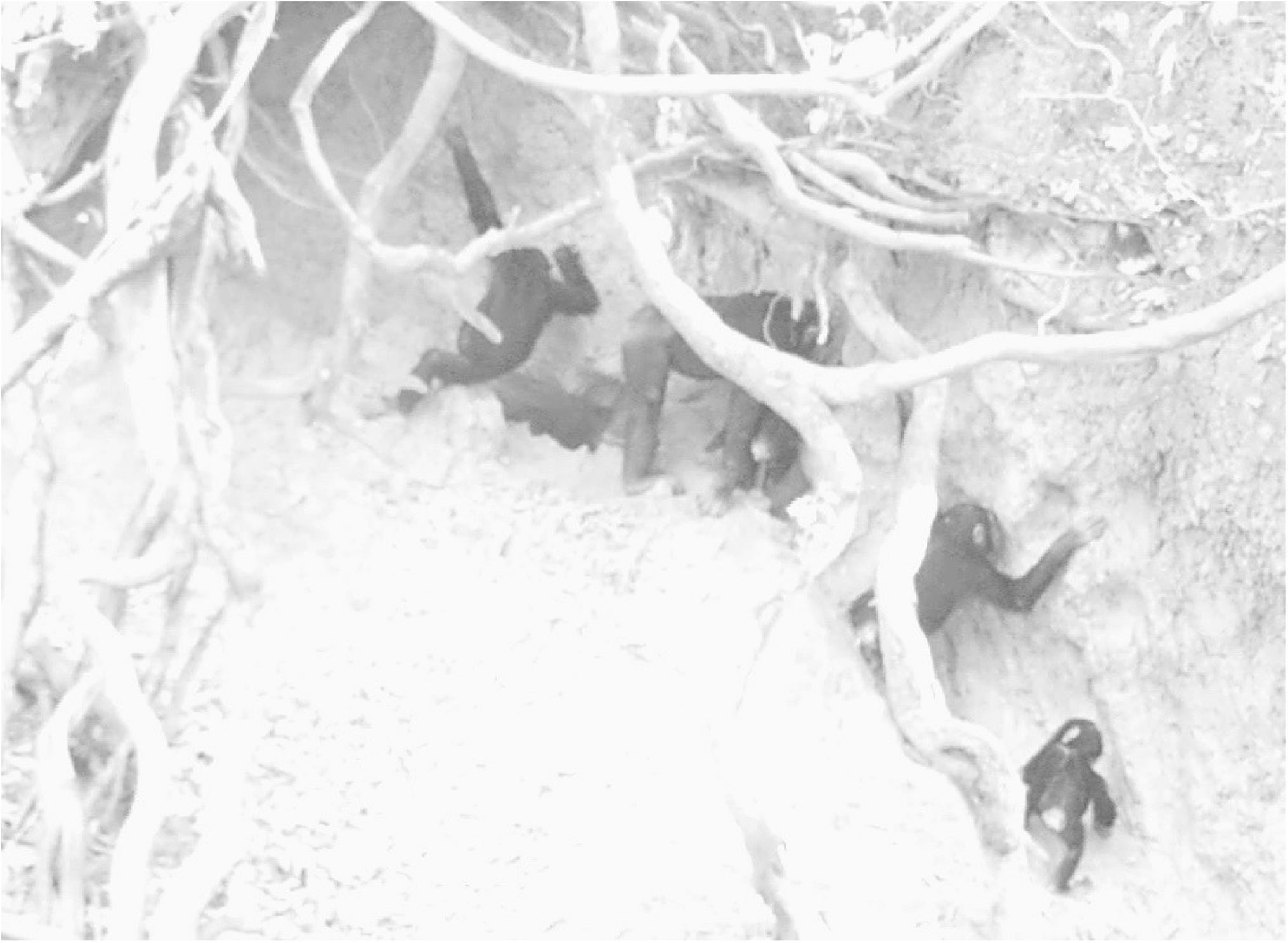
Chimpanzees in the Odyssey community consuming soil in a cliff at the core of their home-range. We can see an adult female, two adolescents and two infants using both teeth and fingernails to scrape the soil. Video still, February 2019.

## SUPPLEMENTARY VIDEO

https://www.youtube.com/watch?v=nyRhqfdGI10&t=104s

## STATEMENTS

This research was carried out following all the international ethical guidelines, the laws of Ivory Coast and with all the permissions to work in a protected area, avoiding any harm or threat to the animals involved. The authors have no conflict of interests. This research was funded by AF, USFWS, FBZ, and had the collaboration of OIPR.

Juan Lapuente designed the study, collected and analyzed the data and wrote the manuscript. K. Eduard Linsenmair contributed to the study design, the interpretation of the results and the manuscript writing. All data generated or analyzed during this study are included in this article. Further inquiries can be directed to the corresponding author.

